# Structural basis of archaeal FttA-dependent transcription termination

**DOI:** 10.1101/2023.08.09.552649

**Authors:** Chengyuan Wang, Vadim Molodtsov, Travis J. Sanders, Craig J. Marshall, Emre Firlar, Jason T. Kaelber, Thomas J. Santangelo, Richard H. Ebright

## Abstract

The ribonuclease FttA mediates factor-dependent transcription termination in archaea^1–3^. Here, we report the structure of a *Thermococcus kodakarensis* transcription pre-termination complex comprising FttA, Spt4, Spt5, and a transcription elongation complex (TEC). The structure shows that FttA interacts with the TEC in a manner that enables RNA to proceed directly from the TEC RNA-exit channel to the FttA catalytic center and that enables endonucleolytic cleavage of RNA by FttA, followed by 5’→3’ exonucleolytic cleavage of RNA by FttA and concomitant 5’→3’ translocation of FttA on RNA, to apply mechanical force to the TEC and trigger termination. The structure further reveals that Spt5 bridges FttA and the TEC, explaining how Spt5 stimulates FttA-dependent termination. The results reveal functional analogy between bacterial and archaeal factor-dependent termination, reveal functional homology between archaeal and eukaryotic factor-dependent termination, and reveal fundamental mechanistic similarities in factor-dependent termination in the three domains of life: bacterial, archaeal, and eukaryotic.

**One sentence summary:** Cryo-EM reveals the structure of the archaeal FttA pre-termination complex

## Main

FttA (also known as aCPSF1) mediates factor-dependent transcription termination in archaea^1–3^, and is responsible for transcription termination at the majority of transcription units in archaea *in vivo*^3^. FttA exhibits endoribonucleolytic activity and 5’→3’ exoribonucleolytic activity^1–7^. FttA has a bi-metal nucleolytic catalytic center, comprising two Zn^2+^ ions coordinated by six histidine residues, located in an active-center groove formed by the interface between a metallo-β-lactamase domain (MβL) and a fused β-CASPase domain (β-CASP)^4–7^. FttA binds specifically to U-rich RNA^3^ and, as a result, performs endoribonucleolytic and exoribonucleolytic activity preferentially on U-rich RNA^2–3,7^. FttA is a homolog of the RNA-cleavage subunits of the multisubunit, megadalton complexes that mediate factor-dependent transcription termination in eukaryotes (Fig. 1a)^1–15^. Thus, FttA is a homolog of INTS11 ,the RNA-cleavage subunit of the eukaryotic integrator complex (INT), a 14-subunit, 1.6 MDa complex involved in termination of Pol II on snRNA genes and promoter-proximally paused Pol II on mRNA genes^8–11^. FttA also is a homolog of CPSF73 (Yth1 in yeast), the RNA-cleavage subunit of the eukaryotic cleavage and polyadenylation factor (CPSF; CPF in yeast), a 21-subunit, 1.8 MDa complex involved in termination of Pol II on mRNA genes^12–15^. Archaeal FttA consists of fused MβL and β-CASP domains preceded by two K homology domains (KH1 and KH2) (Fig. 1a, line 1).^4–7^ Eukaryotic INTS11 and CPSF73 consist of fused MβL and β-CASP domains followed by C-terminal dimerization sequences (Fig. 1a, lines 2-3).^4–9,12,15^

**Fig. 1.**
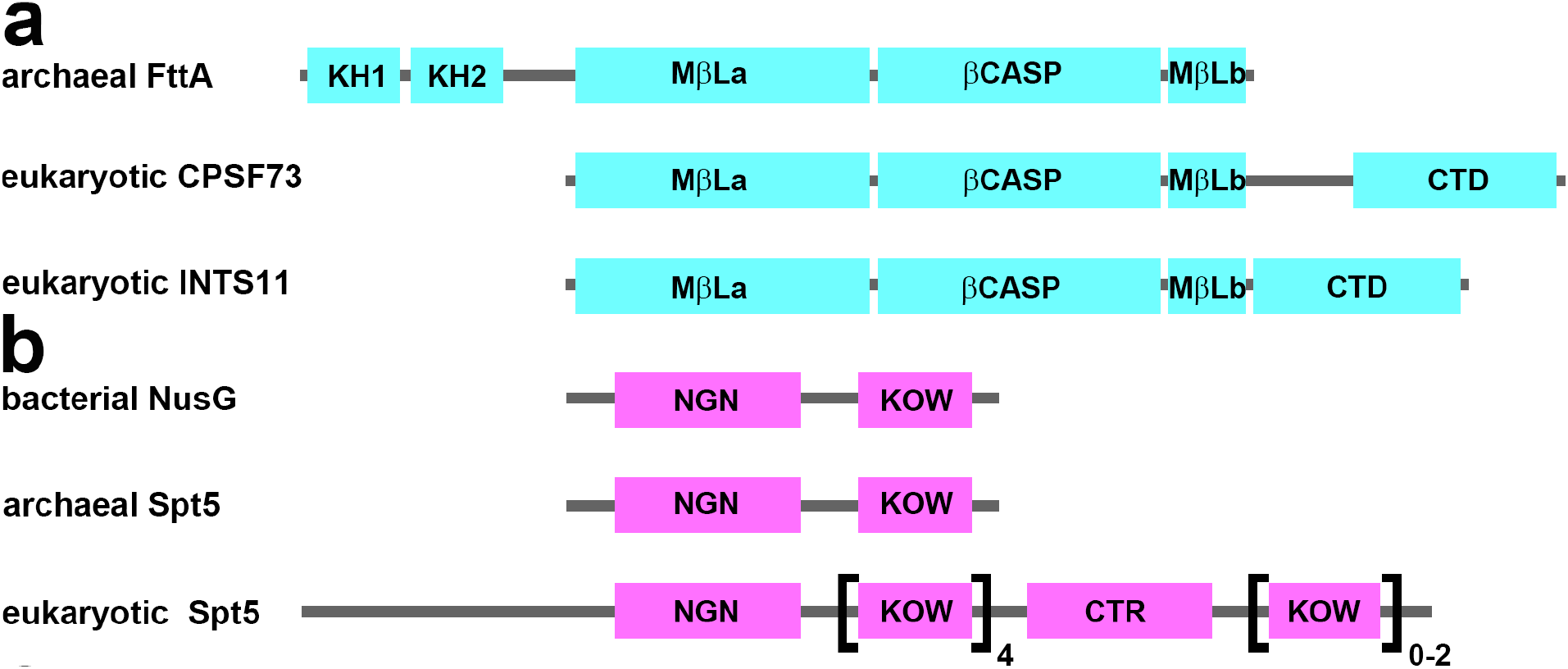
Transcription termination factor FttA and transcription elongation factor NusG/Spt5. (a) Domain architectures of archaeal FttA, eukaryotic CPSF73, and eukaryotic INTS11^4–9,12,15^. (b) Domain architectures of bacterial NusG, archaeal Spt5, and eukaryotic Spt5^16–18^. KOW1-KOW5 are present in all eukaryotes; KOW6-KOW7 are present only in plants and metazoans.

NusG/Spt5 is the sole universally conserved transcription factor, present in bacteria, archaea, and eukaryotes^16–18^. Bacterial NusG and archaeal Spt5 each consist of a NusG N-terminal domain (NGN), which interacts with RNA polymerase (RNAP), and a Kyprides-Ouzounis-Woese domain (KOW), which interacts with factors that associate with and/or cooperate with RNAP (Fig. 1b, lines 1 and 2)^16–18^. Eukaryotic Spt5 consists of an NGN domain, linker and KOW domain followed by eukaryotic-specific C-terminal sequences (Fig. 1b, line 3)^16–18^. FttA-dependent transcription termination is stimulated by Spt5^1^. A protein fragment corresponding to the Spt5 KOW domain is unable to stimulate FttA-dependent termination^1^, indicating that stimulation of FttA-dependent termination requires the Spt5 NGN domain or requires both the Spt5 NGN domain and Spt5 KOW domain^1^.

Bacterial factor-dependent transcription termination is stimulated by NusG, which bridges the transcription elongation complex (TEC) and the transcription termination factor Rho, with the NusG NGN domain interacting with the TEC, and with the NusG KOW domain interacting with Rho^19^. The fact that both bacterial Rho-dependent transcription termination and archaeal FttA-dependent transcription termination are stimulated by a member of the NusG/Spt5 protein family^1,19^ raises the possibility that bacterial Rho-dependent transcription termination and archaeal FttA-dependent transcription termination share functional homology or functional analogy. Here, to test this possibility, we used cryogenic electron microscopy (cryo-EM) to determine an atomic structure of an archaeal FttA-dependent pre-termination complex.

### Structure of archaeal FttA pre-termination complex

To obtain an archaeal FttA pre-termination complex, we analyzed a mutant FttA derivative, FttA^H255A^, from the hyperthermophilic archaeon *Thermococcus kodakarensis* (*Tko*). FttA^H255A^ has a substitution of one of the six histidine residues that coordinate Zn^2+^ ions in the FttA nucleolytic catalytic center^1,4–7^. The FttA^H255A^ substitution reduces, but does not eliminate, endonucleolytic cleavage of RNA and 5’→3’ exonucleolytic cleavage of RNA^1^. FttA^H255A^ is proficient in forming a pre-termination complex that contain both FttA and a TEC, but is deficient in subsequent steps of termination^1^.

We assembled an FttA pre-termination complex from *Tko* FttA^H255A^, *Tko* Spt5, *Tko* Spt4 (the binding partner of Spt5)^16–18^, *Tko* RNAP, and a synthetic nucleic-acid scaffold that contained a 24 nt U-rich 5’ extension of the RNA oligomer able to serve as an FttA binding site (scaffold sequence in Fig. S1). We then determined a 3.9 Å resolution structure of the complex by use of single-particle-reconstruction cryo-EM (FttA^H255A^-Spt4-Spt5-TEC; Figs. 2a and S2; Table S1).

**Fig. 2.**
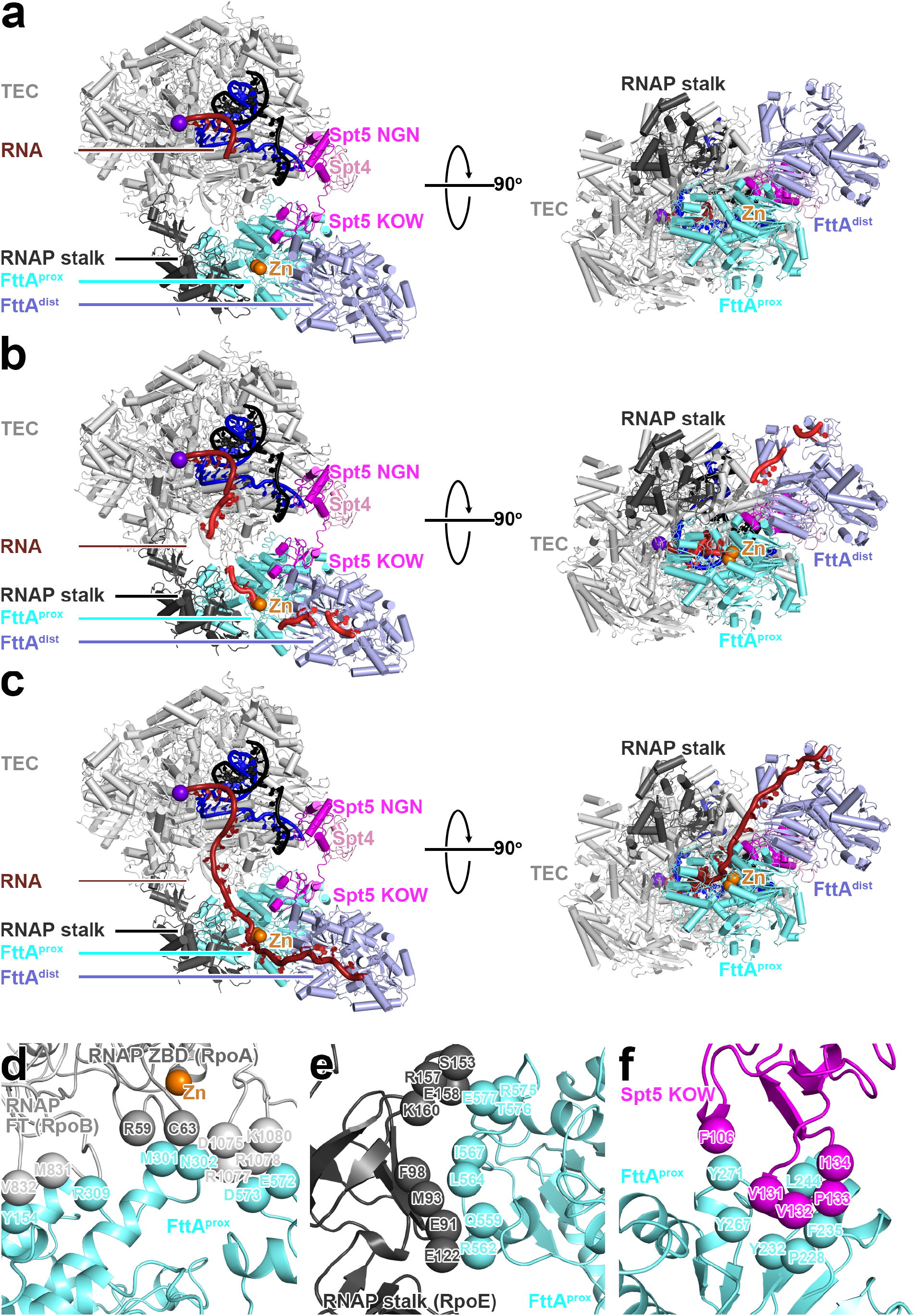
Structure of the FttA pre-termination complex (*Tko* FttA^H255A^-Spt4-Spt5-TEC). (a) Cryo-EM structure. Left panels, view orientation having TEC RNA-exit channel aligned with y-axis, showing passage of RNA through TEC RNA-exit channel and across FttA^prox^ and FttA^dist^. Right panels, orthogonal view orientation showing inferred RNA path across FttA^prox^ and FttA^dist^. FttA^prox^, cyan with nucleolytic active-center Zn^2+^ ions as orange spheres; FttA^dist^, light blue; Spt4, pink; Spt5, magenta; RNAP TEC, gray; RNAP stalk, dark gray; nontemplate-strand DNA, template-strand DNA, and RNA, black, blue, and brick-red, respectively. (b) Cryo-EM structure with superimposed RNA nucleotides in and immediately adjacent to TEC RNA-exit channel (from PDB 7XN7)^21^, RNA nucleotides interacting with FttA^prox^ MβL and β-CASP domains (from PDB 3IE1), and RNA nucleotides interacting with FttA^dist^ KH1 and KH2 domains (from PDB 3AF6 and PDB 3IEV)^5,22^. (c) Cryo-EM structure with inferred full path of RNA through TEC RNA-exist channel, FttA^prox^, and FttA^dist^. (d) Protein-protein interactions between FttA^prox^ and RNAP zinc binding domain 1 (RpoA ZBD1), RNAP zinc binding domain 3 (RpoB ZBD3), and RNAP flap tip (RpoB FT). (e) Protein-protein interactions between FttA^prox^ and RNAP stalk (RpoE). (f) Protein-protein interactions between FttA^prox^ and Spt5 KOW.

The structure shows that a *Tko* FttA dimer (comprising protomer FttA^prox^ located proximal to the TEC and protomer FttA^dist^ located distal to the TEC) interacts with the *Tko* TEC at the mouth of the TEC RNA-exit channel, enabling interactions of the FttA dimer with RNA emerging from the RNA-exit channel (Fig. 2a). The structure further shows that *Tko* Spt5 bridges the TEC and FttA,, with the Spt5 NGN domain interacting with its binding partner Spt4 and the TEC, and with the Spt5 KOW domain interacting with FttA^prox^ (Fig 2a).

The structure shows RNA nucleotides in the TEC RNA-DNA hybrid, but does not show RNA nucleotides in the TEC RNA-exit channel or RNA nucleotides outside the TEC (Fig. 2a). We infer that, during preparation of the sample, endonucleolytic cleavage of the RNA by the mutant FttA derivative occurred, followed by dissociation of the 5’ RNA cleavage fragment and either (i) limited 5’→3’ exonucleolytic cleavage by FttA of the 5’ end of the 3’ RNA cleavage fragment, or (ii) retraction of the 5’ end of the 3’ RNA cleavage fragment into the TEC RNA-exit channel and compaction and disorder of the 5’ end of RNA in the RNA-exit channel (as observed previously in structures having ≥8 nt of RNA in the RNA-exit channel)^20^.

Although RNA nucleotides in the TEC RNA-exit channel and outside the TEC are not resolved in the structure, we are able to model those RNA nucleotides by superimposing published structures of (i) RNA in a TEC RNA-exit channel (PDB 7XN7)^21^, (ii) RNA bound to MβL and β-CASP domains (PDB 3IE1), and (iii) RNA bound to KH1 and KH2 domains (PDB 3AF6 and PDB 3IEV)^5,22^ (Fig. 2b; see refs. 6-7). Connection of the superimposed RNAs, interpolating 3 nt between the modeled RNA segment in the TEC RNA-exit channel and the modeled RNA segment bound to FttA^prox^, interpolating 5 nt between the modeled RNA segment bound to FttA^prox^ and the modeled RNA segments bound to FttA^dist^, and interpolating 1 nt between the modeled RNA segments bound to FttA^dist^, provides a model of the full path of RNA (Fig. 2c).

In the resulting model of the full path of RNA, the RNA proceeds directly from the RNA-exit channel, through the FttA^prox^ active-center groove, to the FttA^prox^ nucleolytic active center (Fig. 2c). RNA then proceeds from the FttA^prox^ nucleolytic active center, through a continuation of the FttA^prox^ active-center groove, to the FttA^dist^ KH1 and KH2 domains (Fig. 2c). 6 nt of RNA, positioned 16-21 nt from the RNA 3’ end in a TEC in the post-translocated state (17-22 nt in the pre-translocated state), interact with FttA^prox^, and 9 nt of RNA, positioned 24-32 nt from the RNA 3’ end in a TEC in the post-translocated state (25-33 nt in the pre-translocated-state), interact with FttA^dist^ (Fig. 2c). The RNA phosphodiester bond 19-20 nt from the RNA 3’ end in a TEC in the post-translocated state (20-21 nt in the pre-translocated state) interacts with the FttA^prox^ nucleolytic active center (Fig. 2c), suggesting that endonucleolytic cleavage by the FttA^prox^ nucleolytic active center occurs 19-20 nt from the RNA 3’ end in a TEC in the post-translocated state (20-21 nt the TEC pre-translocated state). Consistent with this inference, addition of FttA to a TEC has been shown to cleave RNA at a position ∼20-30 nt from the RNA 3’ end^1^.

The RNA interacts with the opposite faces of FttA^prox^ and FttA^dist^, interacting with the face of FttA^prox^ that contains the active-center groove and the nucleolytic catalytic center, and interacting with the face of FttA^dist^ that does not contain the active-center groove and the nucleolytic catalytic center, and, instead, contains the KH1 and KH2 domains (Fig. 2c). We infer that FttA^prox^ and FttA^dist^ play distinct functional roles. FttA^prox^ plays the key active role, carrying out endonucleolytic and endonucleolytic cleavage that drive termination, and FttA^dist^ plays a supporting role, providing an extension of the protein-RNA interaction surface that enables additional interactions with the RNA product and an enhanced stability of association with the pre-termination complex. The inferred involvement of both protomers of the FttA dimer in FttA-RNA interactions in the FttA pre-termination complex is consistent with, and accounts for, the observation that dimerization of FttA enhances the RNA-cleavage activity of FttA^3,7^.

The protein-protein interface that connects TEC and Spt5 to FttA^prox^ is extensive (1,772 Å^2^ buried surface area), suggesting that the translational and rotational orientation of TEC and Spt5 relative to FttA^prox^ is fixed or nearly fixed, and likely remains fixed or nearly fixed, during nucleolytic cleavage and termination by FttA^prox^ . The protein-protein interface has three parts: (i) a set of interactions between the largest and second-largest subunits of RNAP (RpoA and RpoB) and FttA^prox^, comprising interactions by RpoA zinc-binding domain 1 (ZBD1), RpoB zinc-binding domain 3 (ZBD3), and the RpoB flap tip (FT) (Fig. 2d; total buried surface area = 839 Å^2^); (ii) an interaction between the RNAP stalk (RpoE) and FttA^prox^ (Fig. 2e; 395 Å^2^ buried surface area; and (iii) an interaction between Spt5 KOW and FttA^prox^ (Fig. 2f; 538 Å^2^ buried surface area). The involvement of the RNAP stalk and Spt5 in the TEC-FttA interface is consistent with, and accounts for, the observations that the RNAP stalk and Spt5 are essential for FttA-dependent termination^1^.

The dimerization interface between FttA^prox^ and FttA^dist^ also is extensive (Fig. S3; 873 Å^2^ buried surface area). The dimerization interface observed in our structure of the FttA pre-termination complex is identical to the dimerization interface observed in crystal structures of FttA in the absence of other factors^6–7^.

### Mechanism of archaeal FttA-dependent termination

The orientation of FttA^prox^ relative to the TEC in the structure (Fig. 2c), and the position and orientation of FttA^prox^ nucleolytic catalytic site relative to the TEC in the structure (Fig. 2c), support the proposal^1–3^ that FttA mediates termination by performing endonucleolytic cleavage of the RNA followed by processive 5’→3’ exonucleolytic cleavage of the RNA, resulting in 5’→3’ translocation of FttA on the RNA toward the TEC, application of mechanical force to the TEC, disruption of the TEC, and termination. The key premise of this proposed is that each step of processive 5’→3’ exonucleolytic activity by FttA necessarily results in 5’→3’ translocation of FttA relative to RNA, and, in the context of the extensive and likely fixed or nearly fixed protein-protein interface between FttA and the TEC, necessarily applies mechanical force, through the RNA, to the TEC

### Mechanisms of bacterial, archaeal, and eukaryotic factor-dependent termination

Our structure of the archaeal FttA pre-termination complex defines the relationship of archaeal factor-dependent termination to bacterial and eukaryotic factor-dependent termination.

The relationship of archaeal FttA-dependent termination to bacterial Rho-dependent termination is a relationship of *analogy in the absence of homology*. FttA bears no sequence or structural relationships with Rho (Fig. 3a). However, FttA^prox^ interacts with the mouth of the archaeal TEC RNA-exit channel, enabling direct delivery of RNA to the FttA^prox^ catalytic center in a manner that is strikingly similar to the manner in which Rho interacts with the mouth of the bacterial TEC RNA-exit channel, enabling direct delivery of RNA to the Rho catalytic center (Fig. 3a). In addition, Spt5 bridges FttA^prox^ and the archaeal TEC in a manner strikingly similar to manner in which the bacterial Spt5 homolog, NusG, bridges Rho and the bacterial TEC (Fig. 3a).

**Fig. 3:**
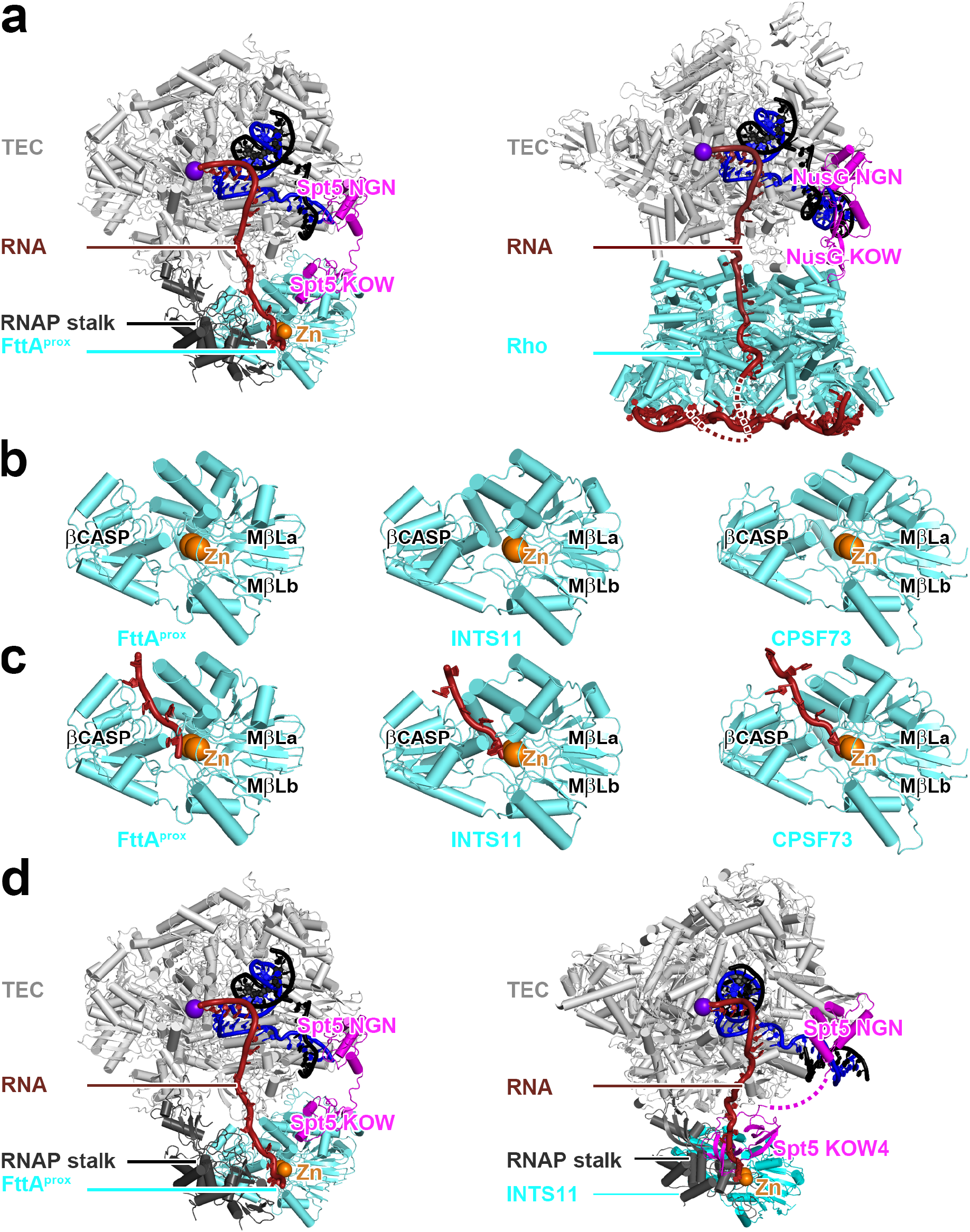
Structural analogy of factor-dependent pre-termination complexes in bacteria and archaea; structural homology of factor-dependent pre-termination complexes in archaea and eukaryotes.. (a) Comparison of archaeal FttA-dependent pre-termination complex (left; Fig. 2) and bacterial Rho-dependent pre-termination complex (right; PDB 8E6W and PDB 8E6X)^19^. In left panel, FttA^prox^ KH1-KH2 domains, FttA^dist^, and Spt4 are omitted for clarity. Rho, cyan; NusG, magenta; Mg-ADP-BeF3 bound to Rho, yellow-orange. Other colors as in Fig 2a. (b) Comparison of MβL and β-CASP domains of archaeal FttA^prox^ (Fig 2), eukaryotic INTS11 (PDB 7CYX)^9^, and eukaryotic CPSF73 (PDB 6V4X)^23^. MβL and β-CASP, cyan; nucleolytic active-center Zn2+ ions, orange spheres. (c) Comparison of RNA-bound MβL and β-CASP domains of FttA^prox^ (Fig. 2), INTS11 (PDB 7CYX)^9^, and CPSF73 (PDB 6V4X)^23^. RNA sgement 3’ to nucleolytic active-center, red. Other colors as in b. (d) Comparison of archaeal FttA-dependent pre-termination complex (left; Fig. 2; FttA^prox^ KH1-KH2 domains, FttA^dist^, and Spt4 omitted for clarity) and eukaryotic INT-dependent pre-termination complex (right; PDB 7YCX)^9^. In left panel, FttA^prox^ KH1-KH2 domains, FttA^dist^, and Spt4 are omitted for clarity. In right panel, INT and PPA2 subunits other than INTS11, INTS11 CTD, RPB1 CTD, NELF, Spt4, and Spt5 KOW domains other than NTD and KOW4 are omitted for clarity. INTS11, cyan; connector between SPT5 NGN and Spt5 KOW4, dashed magenta line. Other colors as in Fig. 2a.

The relationship of archaeal FttA-dependent termination to eukaryotic factor-dependent termination is one of *homology*. The structures of the MβL and β-CASP domains of FttA^prox^ in the FttA pre-termination complex are superimposable on the structures of the MβL and β-CASP domains of eukaryotic INTS11^8–9^, the RNA cleavage subunit of the 14-subunit, 1.6 MDa INT complex^8–11^, and of eukaryotic CPSF73^23^, the RNA cleavage subunit of a 21-subunit, 1.8 MDa complex CPSF complex^12–15^ (Fig. 3b). In addition, the inferred protein-RNA interactions made by the MβL and β-CASP domains of FttA^prox^ in the FttA pre-termination complex are superimposable on those made by eukaryotic INTS11 and eukaryotic CPSF73 (Fig. 3c)^8–9,23^. Most important, the translational and rotational orientation of archaeal FttA^prox^ relative to the archaeal TEC and Spt5 KOW are closely similar to those of eukaryotic INTS11 and Spt5 KOW4 in published structures of the 14-subunit, 1.6 MDa human INT complex and Spt5 bound to human Pol II (Fig. 3d)^8–9^.

Fig. 4 summarizes the mechanisms of factor-dependent termination in the three domains of life: bacteria, archaea, and eukaryotes. In bacterial factor-dependent termination, the RNA translocase Rho fbinds to RNA emerging from a TEC, stabilized by a bridging interaction by NusG,, yielding a pre-termination complex (Fig. 4a, left); Rho then performs ATP-hydrolysis-dependent 5’→3’ translocation on the RNA, thereby applying mechanical force to the TEC and triggering termination (Fig. 4a, right)^19^. In archaeal FttA-dependent transcription termination the ribonuclease FttA binds to RNA emerging from a TEC, stabilized by a bridging interaction by Spt5, and FttA performs endonucleolytic cleavage and limited 5’→3’ exonucleolytic cleavage of the RNA, yielding a pre-termination complex (Fig. 4b, left); FttA then performs processive 5’→3’ exonucleolytic cleavage of the RNA, and concomitant 5’→3’ translocation on the RNA, thereby applying mechanical force on TEC and triggering termination (Fig. 4b, right; this work). In eukaryotic INT- or CPSF-dependent transcription termination, an INTS11- or CPSF73-containing multisubunit, megadalton complex binds to RNA emerging from a TEC, stabilized by a bridging interaction by Spt5, and INTS11 or CPSF73 performs endonucleolytic cleavage and limited 5’-3’ exonucleolytic cleavage of the RNA, forming a pre-termination complex (Fig. 4c, left); then either the INTS11- or CPSF73-containing complex, or an Xrn2-containing complex (Rat1-containing complex in yeast), performs processive 5’→3’ exonucleolytic cleavage of the RNA, and concomitant 5’→3’ translocation on the RNA, thereby applying mechanical force to the TEC and triggering termination (Fig. 4c, right)^8–15,24–30^. Available evidence suggests that the mechanism shown in the upper right subpanel of Fig. 4c predominates for INT-mediated termination^13,29^, and the mechanism in the lower right subpanel of Fig. 4c predominates for CPSF-mediated termination^12–15,24–30^.

**Fig. 4:**
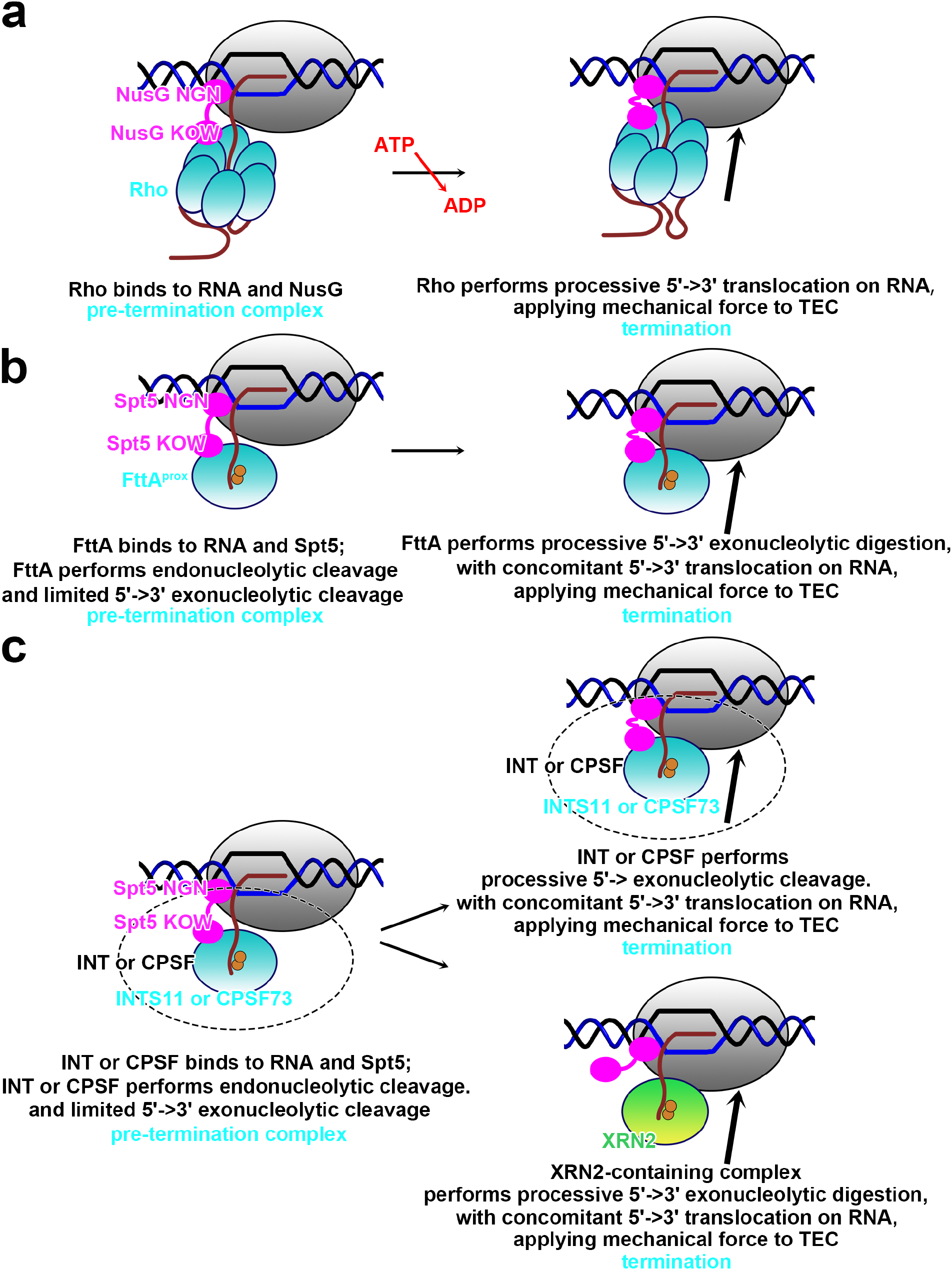
Mechanisms of bacterial, archaeal, and eukaryotic factor-dependent termination. (a) Bacterial Rho-dependent termination^19^. (b) Archaeal FttA-dependent termination (Figs. 2-3). (c) Eukaryotic INT-dependent termination (INTS11 as FttA homolog) and CPSF-dependent termination (CPSF73 as FttA homolog)^8–15,24–30^. The upper-right pathway is thought to predominates for INT-dependent termination^13–29^; the lower-right pathway is thought to predominate for CPSF-dependent termination^12–15,24–30^.

The mechanisms proposed in Fig. 4 suggest a fundamental mechanistic unity of factor-dependent termination across the three domains of life: bacteria, archaea, and eukaryotes. *In each case, factor-dependent termination is proposed to involve 5’→3’ translocation on RNA by a termination factor that interacts with RNA emerging from the TEC RNA-exit channel, 5’→3’ translocation on RNA by the termination factor is proposed to apply mechanical force to the TEC, and the application of mechanical force to the TEC is proposed to triggers termination.* Consistent with these proposals, single-molecule force measurements show that the application of mechanical force through RNA to a TEC can trigger termination^31^. The specific reactions through which the application of mechanical force through RNA,to a TEC triggers termination, inhibiting RNA extension and predisposing the TEC to dissociation, remain to be elucidated, It is likely that these reactions include one or more of: (i) forward translocation of the TEC without nucleotide addition (hypertranslocation model)^31–35^, (ii) extraction of RNA from the TEC (RNA extraction model)^31,34–35^, and (iii) reorganization of TEC structure (allosteric model)^34–37^.

## Discussion

In bacteria and archaea, there is no nuclear membrane, and this transcription and translation occur in the same cellular compartment and can be physically coupled^20;38–41^. In bacteria, Rho not only mediates factor-dependent transcription termination, but also mediates quality control of transcription-translation coupling, detecting lapses in transcription-translation coupling, such as losses of transcription-translation coupling during translational arrest, and resolving lapses in transcription-translation coupling through transcription termination^19,,39,41^. The structures of the *E. coli* Rho-dependent pre-termination complex^19^ and the *E. coli* coupled transcription-translation complex^20,38^ indicate that the mechanism of Rho-dependent quality control of transcription-translation coupling relies on a simple steric-exclusion: mechanism: namely, the protein-protein interface of the TEC and NusG with Rho in the Rho pre-termination complex is sterically incompatible with, and mutually exclusive with, the protein-protein interface of the TEC and NusG with the bacterial ribosome in the coupled transcription-translation complex^19–20,38–39^. It is attractive to hypothesize that the mechanism of quality control of transcription-translation coupling in archaea is analogous: namely, a simple steric-exclusion mechanism, based on sterically incompatible, and mutually exclusive, protein-protein interfaces for interactions of the TEC and Spt5 with FttA and interactions of the TEC and Spt5 with the archaeal ribosome. Testing this hypothesis will require the determination of a structure of the archaeal coupled transcription-translation complex and assessment of potential steric incompatibility and mutual exclusivity of the interactions therein with the interactions in the archaeal FttA pre-termination complex.

Our results define the atomic structure of the archaeal FttA pre-termination complex, suggest the likely mechanism of termination, suggest a fundamental unity in the mechanism of factor-dependent termination, in bacteria, archaea, and eukaryotes, and provide a foundation for further structural and functional understanding of archaeal transcription termination and archaeal transcription-translation-coupling quality control.

## Methods

### Proteins

*Tko* FttA^H255A^, *Tko* Spt4, *Tko* Spt5, and *Tko* RNAP were prepared as described^1^. Prior to sample preparation for cryo-EM structure determination, each protein sample was dialyzed overnight at 4°C against 2 L storage buffer (10 mM Tris-HCl, pH 7.5, 100 mM NaCl, 0.1 mM EDTA, and 5 mM dithiothreitol), concentrated by centrifugation at 4,000xg at 4°C through VivaSpin 10 kDa MWCO concentrators (Sartorius), aliquotted, flash-frozen in liquid nitrogen, and stored at -80°C.

### Nucleic acids

Oligodeoxyribonucleotides and oligoribonucleotides (sequences in Fig. S1) were purchased (Integrated DNA Technologies), PAGE-purified, dissolved in annealing buffer (5 mM Tris-HCl, pH 7.5) to 1 mM, and stored at -80°C in aliquots. Nucleic-acid scaffolds were prepared by mixing 100 µM nontemplate-strand oligodeoxyribonucleotide, 100 µM template-strand oligodeoxyribonucleotide, and 100 µM oligoribonucleotide in 100 µl water, heating 10 min at 95°C, and cooling over 3 h to 22°C. The resulting nucleic-acid scaffolds were aliquotted and stored at -80°C.

### Termination assays

Termination assays were performed as described^1^.

### Cryo-EM structure determination: sample preparation

Reaction mixtures containing, 8 µM *Tko* Spt4, 8 µM *Tko* Spt5, 4 µM *Tko* RNAP , and 5 μM nucleic-acid scaffold in 100 μl transcription buffer (20 mM HEPES-NaOH, pH 7.5, 30 mM KCl, 10 mM magnesium acetate, and 1 mM dithiothreitol) were incubated 10 min at 22°C, and then were supplemented with 30 μl 60 µM FttA**^H^**^255^**^A^** and incubated 10 min at 22°C and 5 min at 42°C. Reaction mixtures were concentrated to 35 µl by centrifugation 12 min at 20,000xg at 4°C through pre-chilled Amicon Ultra-0.5 30 KDa MWCO concentrators (EMD Millipore), were supplemented with 4 µl of ice-cold 80 mM CHAPSO (Hampton Research), and were kept on ice until applied to EM grids.

EM grids were prepared using a Vitrobot Mark IV autoplunger (FEI/ThermoFisher), with the environmental chamber set to 22°C and 100% relative humidity. Samples (3 μl) were applied to Quantifoil 2/1 Cu 300 holey-carbon grids (Quantifoil) glow-discharged 60 s using a PELCO glow-discharge system (Ted Pella); grids then were blotted with #595 filter paper (Ted Pella) for 7-8 s, flash-frozen by plunging in liquid ethane cooled with liquid N_2_, clipped, and stored in liquid N_2_.

### Cryo-EM structure determination: data collection and data reduction

Cryo-EM data were collected at the National Center for Cryo-EM Access and Training (NCCAT), using a 300 kV Krios Titan electron microscope (FEI/ThermoFisher) equipped with a Gatan K3 Summit direct electron detector (Gatan). Images were collected in counting mode using Leginon^42^ at a nominal magnification of 81,000x, a calibrated pixel size of 1.069 Å/pixel, and a dose rate of 26.252 electrons/Å^2^/s. Movies were recorded at 50 ms/frame for 2.5 sec (50 frames), resulting in a total radiation dose of 65 electrons/Å^2^. Defocus range varied from -0.8 μm and -2.0 μm. A total of 17,601 micrographs were recorded from one grid over three days. Micrographs were gain-normalized and defect-corrected.

Data were processed as summarized in Fig. S2a. Data processing was performed using a Tensor TS4 Linux GPU workstation with four GTX 1080 Ti graphic cards (NVIDIA). Dose weighting and motion correction (5×5 tiles; b-factor = 150) were performed using Motioncor2^43^. CTF estimation was performed using CTFFIND4^44^. Subsequent image processing was performed using Relion 3.1^45^. Automatic particle picking yielded an initial set of 3,382,400 particles. Particles were binned 2x, extracted into 384×384 pixel boxes, and subjected to six rounds of reference-free 2D classification and removal of poorly populated classes, yielding a selected set of 142,151 particles. An *ab initio* model was generated and was subjected to heterogeneous refinement to sub-classify particles. A class of 111,374 particles exhibiting an intact TEC and a large additional density feature was selected and was subjected to homogeneous refinement following the post-processing with a soft mask, yielding a reconstruction with a global resolution of 3.9 Å as determined from gold-standard Fourier shell correlation (Fig S2d; Table S1). The reconstruction showed clear density for two FttA protomers (FttA^prox^ and FttA^dist^), Spt4, Spt5, and the TEC (Fig. S2e-k).

The initial atomic model for the FttA pre-termination complex was built by manual docking in , using Chimera1.14^46^ of: (i) a cryo-EM structure of *Tko* RNAP (PDB 6KF9)^47^, (ii) a homology model of *Tko* FttA built based on a crystal structure of *Pyrococcus horikoshii* FttA (PDB 3AF5)^5^, and (iii) homology models of *Tko* Spt4, *Tko* Spt5 NGN, and *Tko* Spt5 KOW built based on a crystal structure of a *Pyrococcus furiosus* Spt4-Spt5 complex (PDB 3P8B)^48^. DNA and RNA were manually built using Coot^49^. For the FttA^prox^ N-terminus (residues 1-8 and 51-56), FttA^dist^ N-terminus (residues 1-8 and 51-56), Spt4 N-terminus (residues 1-3), Spt5 N-terminus (residues 1-3), RNAP RpoA N-terminus (residues 1-4), RNAP RpoB N-terminus (residues 1-8), RNAP RpoC N-terminus (residue 1), RpoD N-terminus (residue 1), RpoH N-terminus (residues 1-5), RpoK N-terminus (residue 1), and RpoP N-terminus (residues 1-5), density was absent, suggesting high segmental flexibility; these segments were not fitted.

The initial model was subjected to real-space rigid-body refinement in Phenix^50^, and was further refined using secondary-structure, geometry, Ramachandran, rotamer, Cβ, non-crystallographic-symmetry, and reference-model restraints in Phenix. The Spt5 interdomain linker (residues 85-92), RNAP ZBD 1 (RpoA residues 25-75), RNAP ZBD 3 (RpoB residues 1069-1097), RNAP FT (RpoB residues 890-912), and part of the RNAP stalk (RpoE residues 81-181) were subjected to iterative cycles of model building and refinement in Coot^49^. Structure visualization was performed using PyMOL (Schrödinger).

The final atomic density map and atomic coordinates were deposited in the Protein Data Bank and the Electron Microscopy Data Bank with accession codes 8TEV and EMD-41203, respectively.

## Acknowledgments

We thank the Rutgers CryoEM and Nanoimaging Facility and the National Center for CryoEM Access and Training (supported by NIH grant GM129539, Simons Foundation grant SF349247, and New York state grants) for microscope access and the SIII Center (supported by National Natural Science Foundation of China grant 32270039 to C.W.) for computer resources

## Author contributions

V.M., T.J. Santangelo, and R.H.E. designed experiments. V.M., C.J.M, T.J. Sanders, and T.J. Santangelo prepared proteins and nucleic acids and performed biochemical experiments. V.M, C.W., E.F., and J.T.K. performed cryo-EM data collection. V.M., C.W., T.J. Santangelo, and R.H.E. analyzed data. C.W. and R.H.E. prepared figures. R.H.E. wrote the manuscript.

## Data availability statement

Cryo-EM maps have been deposited in the Electron Microscopy Database (EMDB accession code EMD-41203), and atomic coordinates have been deposited in the Protein Database (PDB accession code 8TEV). Unique biological materials will be made available to qualified investigators on request.

## Supplementary Figure Legends

**Fig. S1.**
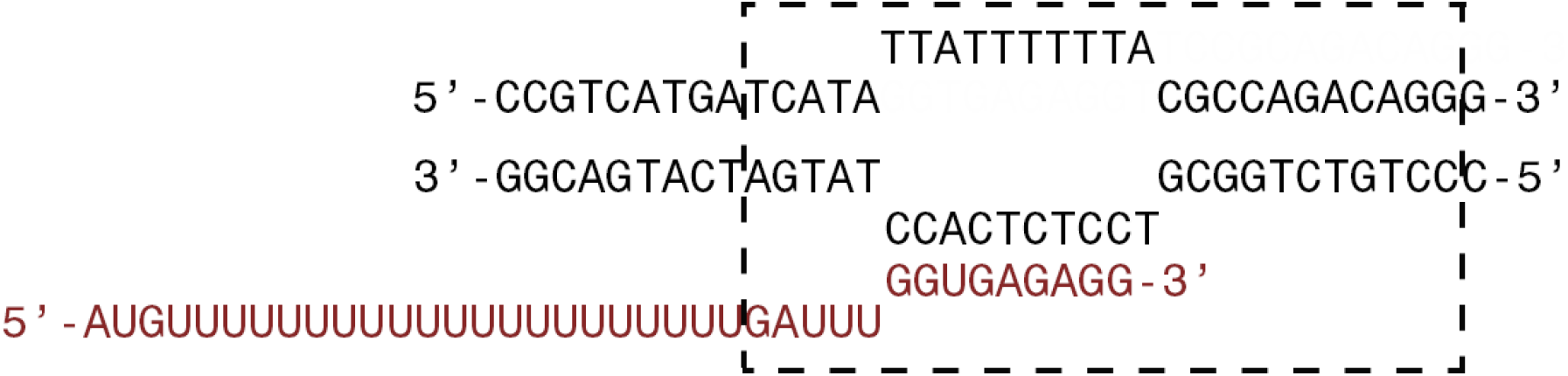
Nucleic-acid scaffold used for structure determination. Black, DNA; brick red, RNA; dashed rectangle, TEC.

**Fig. S2.**
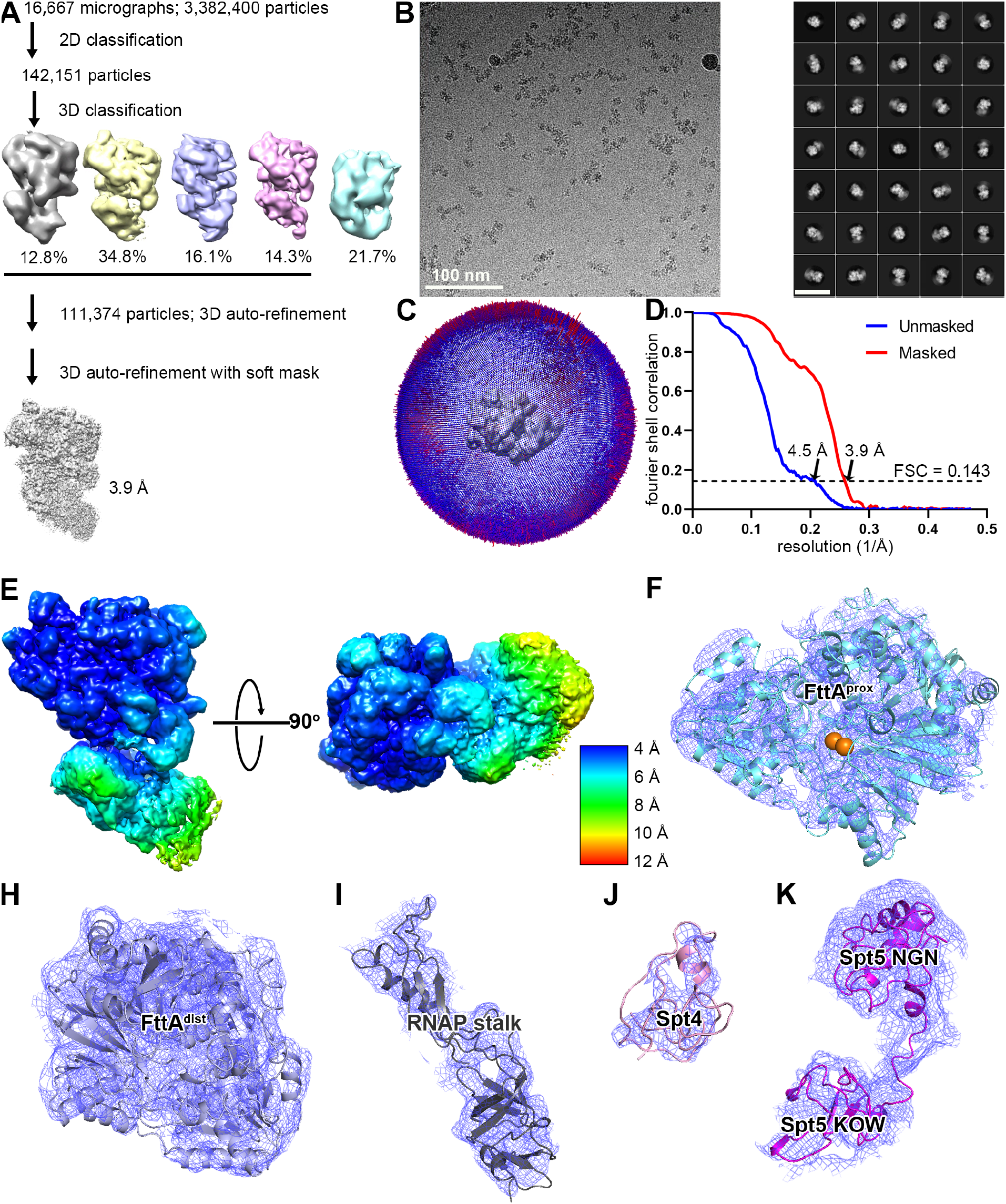
Structure determination: *Tko* FttA^H255A^-Spt4-Spt5-TEC. (a) Data processing scheme (Table S1). (B) Representative electron microphotograph and 2D class averages (50 nm scale bar in right subpanel). (b) Orientation distribution. (c) Fourier-shell correlation (FSC) plot. (e) EM density map colored by local resolution (view orientation as in Fig. 2, left). (f-k) Representative EM density (blue mesh) and fits (ribbons) for FttA^prox^, FttA^dist^, RNAP stalk (RpoE), Spt4, and Spt5.

**Fig S3.**
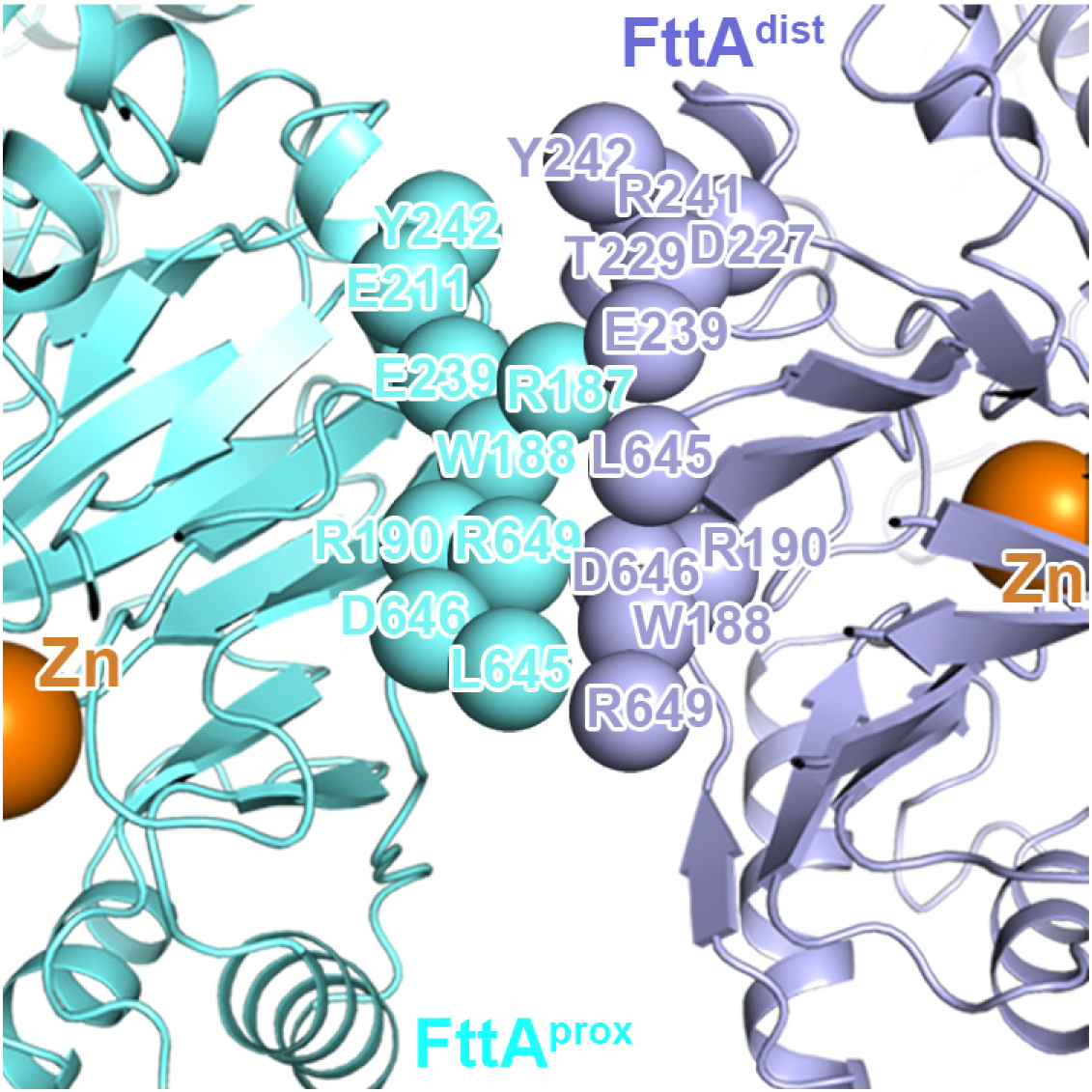
Protein-protein interactions between FttA^prox^ and FttA^dist^.

**Table S1:** Cryo-EM structure: Tko FttA^H255A^-Spt4-Spt5-TEC.

|  | **FttA-Spt4-Spt5-TEC**<br>**(EMDB-41203)**<br>**(PDB 8TEV)** |
| --- | --- |
| **Data collection and processing** |  |
| Magnification | 81,000 |
| Voltage (kV) | 300 |
| Electron exposure (e-/Å<sup>2</sup>) | 65 |
| Defocus range (μm) | -0.8 to -2.0 |
| Pixel size (Å) | 1.069 |
| Symmetry imposed | C1 |
| Initial particle images (no.) | 18,276 |
| Final particle images (no.) | 17,601 |
| Map resolution (Å) | 3.9 |
| FSC threshold | 0.143 |
| Map resolution range (Å) | 3.9-12.2 |
| **Refinement** |  |
| Initial model used (PDB code) | 6KF9, 3AF5, 3P8B |
| Model resolution (Å) | 3.8 |

